# Neural Distinctiveness and Reinstatement of Hippocampal Representations Support Unitization for Associations

**DOI:** 10.1101/2022.07.13.497755

**Authors:** S. Ricupero, C.M. Carpenter, A.C. Steinkrauss, C.R. Gerver, J.D. Chamberlain, R.G. Monkman, A.A. Overman, N.A. Dennis

## Abstract

The medial temporal lobe (MTL) is critical to associative memory success. Yet not all types of associations may be processed in a similar manner within MTL subregions. In particular, work suggests that intra- and inter-item associations not only exhibit differences in overall rates of recollection, but also recruit different MTL subregions. Whereas intra-item associations, akin to unitization, take advantage of associations between within-item features, inter-item associations form links across discrete items. The current work aimed to examine the neural differences between these two types of associations using multivariate neural analyses. Specifically, the current study examined differences across face-occupation as a function of whether the pairing was viewed as a person performing the given job (intra-item binding) or a person saying that they knew someone who had a particular job (inter-item binding). The results show that at encoding, successfully recollected intra- and inter-item associations are discriminable from one another in the hippocampus, parahippocampal cortex, and perirhinal cortex. Additionally, the two trial types are reinstated distinctly such that inter-item trials have higher neural reinstatement from encoding to retrieval compared to intra-item trials in the hippocampus. We conclude that intra- and inter-associative pairs may utilize similar neural regions that represent patterns of activation differentially at encoding. However, in order to reinstate information to the same degree (i.e., subsequently successfully recollected) inter-item associations may act in a compensatory manner, while it is not necessary for intra-item associations to be reinstated to the same degree. This may indicate that intra-item associations promote more efficient reinstatement.

## 1. Introduction

Associative memory allows us to bind together multiple pieces of information from our environment. Examples of everyday associative memory include the binding of faces with their corresponding names and appointments with their corresponding times. Prior research identifies different types of associative binding that depend on the type of information that is bound, such as intra-item and inter-item associations (Ahmad & Hockley, 2014; Bastin et al., 2013; Delhaye et al., 2014; Diana et al., 2007; Giovanello et al., 2006; Parks & Yonelinas, 2015; Parra et al., 2009; Quamme et al., 2007). Intra- and inter-item binding consist of two related, yet unique processes. Inter-item binding is described as the association of across-item features with non-overlapping representations (i.e., an unrelated word pairing like machine-orange; Park & Rugg, 2011), whereas intra-item binding is described as the association of within-item features with overlapping representations (i.e., a related compound word pair like mail-box; Jäger et al., 2006). Intra-item binding aligns with the concept of unitization, in which unique items are bound together in such a way that the resulting ‘unitized’ association is considered a new ensemble that functions similarly to that of a single item within memory, resulting in greater memory accuracy compared to inter-item associations (Ahmad & Hockley, 2014; Bastin et al., 2013; Delhaye et al., 2014; Giovanello et al., 2006; Parks & Yonelinas, 2015; Parra et al., 2009; Quamme et al., 2007). The current study seeks to investigate differences in how these two types of binding operate by examining neural distinctiveness and reinstatement related to the binding and memory for face-occupation associations.

Previous work has used various experimental designs and methodologies to induce intra-item and inter-item associations (Ahmad & Hockley, 2014; Bastin et al., 2013; Overman & Stephens, 2013; Parra et al., 2009; Quamme et al., 2007). For example, intra-item association has been induced by asking participants to formulate a compound word between two unrelated words (e.g., slope-bread), generating a unique definition for that new compound word, which resulted in higher associative memory strength than the unrelated associative condition (Haskins et al., 2008) or by asking participants to use two unrelated words together to create a meaningful sentence (Quamme et al., 2007). The foregoing studies show a memory advantage for intra-compared to inter-item associations. The benefit of these associations is thought to be due to how intra-item, or unitized, word pairs, provide a more holistic and unified representation of the pairing, thus allowing familiarity to support associative memory processes (Ahmad & Hockley, 2014; Bastin et al., 2013; Diana et al., 2011; Quamme et al., 2007).

In addition to a memory advantage for intra-compared to inter-item associations, neuroimaging studies show that the neural correlates of the two types of processing also differ. Specifically, while both types of associative memories have elicited blood-oxygen-level dependent (BOLD) activation within the medial temporal lobe (MTL), including the hippocampus (HC) and parahippocampal cortex (PHC); Allen et al., 2014; Dennis et al., 2014; Piekema et al., 2010; Staresina & Davachi, 2010; Yonelinas et al., 2001), research shows that activation in the perirhinal cortex (PrC) is elicited during the encoding of intra-item unitized pairings (Haskins et al., 2008; Jäger et al., 2006; Staresina & Davachi, 2010), whereas the hippocampus underlies inter-item binding (Dennis, Johnson, et al., 2014; Piekema et al., 2010; Staresina & Davachi, 2010). Such differences in the location and extent of activation across intra- and inter-item associations suggest that the two types of associations are distinct in how they are processed within the MTL.

While the foregoing work focuses on differences in the location and extent of activation using univariate analyses to examine intra- and inter-item associations, the current study sought to extend this work by using multivariate analysis to determine whether the advantage afforded by intra-item associations is related to better discriminability and distinctiveness in the neural patterns of activations for this associative binding. Multivoxel pattern analysis (MVPA) has been used to identify unique neural patterns associated with stimuli of different categories, such as faces and houses (Haxby et al., 2000; Ishai et al., 2000), but also to identify more subtle discrimination across behavior, including true and false memories (Carpenter et al., 2021; Chadwick et al., 2016; Chamberlain et al., 2022), forgotten memories (LaRocque et al., 2013), and recollection and familiarity (Kafkas et al., 2017). Critical to the current study, past work has shown that associative pairs that are similar in content, yet differ with respect to presentation history, are discriminable within MTL subregions, specifically the PHC (Elbich et al., 2021). Prior work has also utilized neural reinstatement and encoding-retrieval similarity analyses to assess how neural patterns are correlated across memory phases, in an effort to investigate overlap in cognitive processes between encoding and retrieval (Hill et al., 2021; Koen, 2022; Koen et al., 2020; Kuhl & Chun, 2014; Ritchey et al., 2013; Thakral et al., 2017). Using this method, work has found that neural pattern reinstatement in MTL regions, including the HC and PHC, has been shown to support associative recollection (Gordon et al., 2014; Staresina et al., 2012). This overlap in representations of patterns from encoding to retrieval has been found to be critical to later successful recollection (see Xue, 2018 for review). We aim to use MVPA and neural reinstatement analyses in the current study to identify any differences in the neural patterns underlying intra- and inter-item associations at each memory stage within the MTL, as well as examine whether the two types of associations show differential neural recapitulation from encoding to retrieval.

Specifcally, we aimed to induce intra-item associations and inter-item associations of face-occupation pairings through the use of different binding strategies. Intra-item associations were created by asking participants to imagine the person as having the identity of the named occupation and imagine the face doing a task related to the occupation. Inter-item associations were created by asking participants to imagine the person speaking aloud that they knew someone with that occupation (see Overman & Stephens, 2013). Based on previous work (Bastin et al., 2013; Delhaye et al., 2014; Diana et al., 2008; Jäger et al., 2006; Quamme et al., 2007), we hypothesize that the ‘doing’ condition would function akin to intra-item associations, demonstrating higher hit rates compared to the ‘speaking,’ or inter-item associations, given that the occupation is stated as a characteristic of the person in the ‘doing’ condition and viewed as separate from the person in the ‘speaking’ condition. Neurally, we expect inter- and intra-associative targets to show neural discriminability within MTL regions related to associative binding, including the HC and PHC as well as in the PrC, as intra-item associations may be unitized as an integrated representation (Jäger et al., 2006; Staresina & Davachi, 2010). With respect to neural distinctiveness, we predicted that intra-item pairs would show greater distinctiveness than inter-item pairs, (Bastin et al., 2013; Delhaye et al., 2014; Diana et al., 2008; Jäger et al., 2006; Quamme et al., 2007) and thus, greater discriminability, compared to inter-item pairs specifically in the PrC and HC based on previous neuroimaging work in unitization that suggests different association types rely on different neural processes (Jäger et al., 2006; Staresina & Davachi, 2010). Finally, with respect to neural reinstatement, we predicted that successfully recollected inter-item pairs would have higher reinstatement compared to intra-item pairs, due to the greater difficulty inherent in remembering inter-item pairs (Ahmad & Hockley, 2014; Bastin et al., 2013; Diana et al., 2011; Parks & Yonelinas, 2015), thus requiring greater strength in neural reinstatement when successful.

## 2. Materials and Methods

### 2.1 Subjects

28 younger adults were recruited from The Pennsylvania State University. Participants were screened for history of psychiatric and neurological illness, head injury, stroke, learning disability, medication that affects cognitive and physiological function, and substance abuse. On the day of the study, all participants provided written informed consent for a protocol approved by The Pennsylvania State University institutional review board. All participants were native English speakers or had learned English before the age of 8, with normal or corrected-to-normal vision and were right-handed. All participants were enrolled in college or postgraduate education. All 28 younger adults were included in all analyses (*M*_age_= 22.11, *SD*_age_= 0.57, range = 18-28; 24 female, 4 male). Participants identified as white (n = 15), as well as Asian/Pacific Islander (n = 10), and more than one race (n = 3), and were all well-educated (*M*years = 15, *SD*years = 0.36).

### 2.2 Regions of Interest (ROIs)

Based upon previous work mentioned above, we restricted our analysis to MTL subregions, including the bilateral perirhinal cortex (PrC), the bilateral parahippocampal cortex (PHC) and the bilateral hippocampus (HC). The ROIs were defined anatomically and created using the human AAL Pickatlas through SPM12 (Lancaster et al., 2000).

### 2.3 Stimuli & Procedure

The current design and stimuli were modified from Overman & Stephens, 2013. The experimental stimuli consisted of 144 black and white photographs of faces (see Criss & Shiffrin, 2004 for standardization detials) and 144 single-word occupations (Ex: pianist; welder), with a majority of the occupations taken from Yovel & Paller (2004) and additional occupations added as needed. During encoding, participants were shown an image of a face and a name tag stating “Hello, I’m [occupation]” or a speech bubble stating “I know [occupation]” (See figure 1). Participants were asked to imagine the face-occupation association and remember the pairings using one of two strategies designed to promote either intra- or inter-item binding. Specifically, the ‘doing’ condition described above was utilized to promote unitization or intra-item binding wherein the participants were asked to imagine the pictured individual performing actions related to the occupation. The ‘speaking’ condition described above was utilized to promote inter-item binding wherein the participants were asked to imagine the pictured individual knowing someone else with the given occupation. Participants were prompted by an instruction screen as to which strategy they should be using, with both encoding strategies used in each run. Participants were also asked to indicate, using a four-point button box, how easy or how difficult it is to use the given strategy for each unique face and job pairing (response options included: very difficult, somewhat difficult, somewhat easy, very easy). Finally, the background screen color randomly alternated between either yellow or blue, with each appearing 50% of the time. (Analyses related to this color manipulation were not included in the current set of analyses).

**Figure 1.**
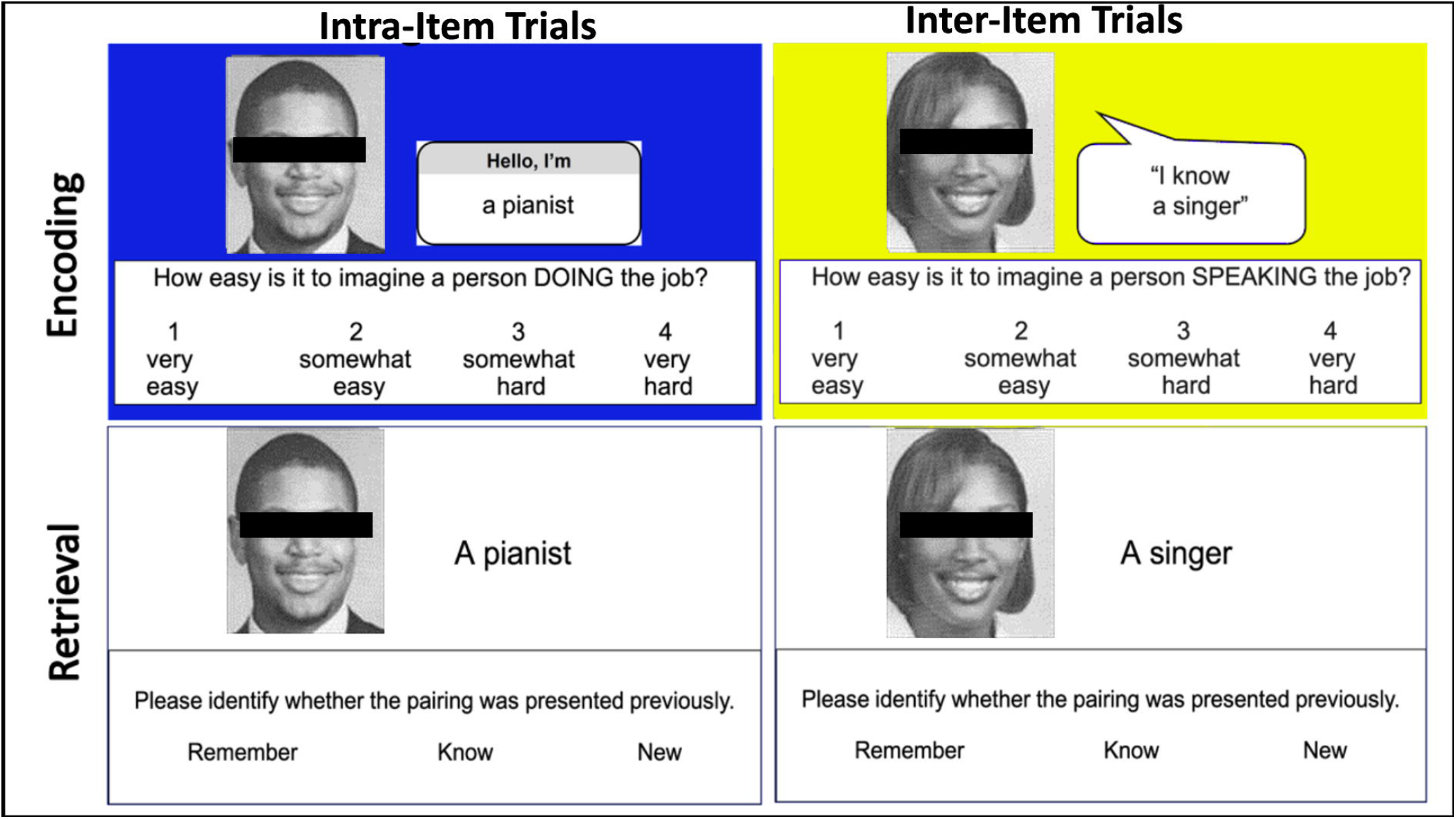
Example stimuli for the associative memory task. Face and job pairings were presented on one of two background colors (yellow or blue) as an incidental feature of the encoding task. Both the encoding conditions contained blue and yellow trials. At retrieval, the pairings were presented with a white background color.

During encoding trials, participants received a prompt (2.5 seconds) to use one of the strategies (i.e., speaking) and saw 9 trials then a prompt to use the other strategy (i.e., doing) and saw 9 trials and this process repeated itself twice per run^1^. During retrieval trials, participants were presented with both target pairs and recombined lure pairs on a white background. Most face job pairs were recombined within the same condition^2^. Participants responded during retrieval using the standard Remember-Know-New paradigm (Yonelinas, 2002). They responded ‘remember’ if they remembered specific details about the face-occupation pair, ‘know’ if they believed they had seen the pairs together previously, but could not remember specific details of the pair, and ‘new’ if they believed the pair was not presented together previously. Each retrieval run included 36 trials, 24 targets and 12 recombined lures. The order of runs included two encoding runs, followed by two retrieval runs, repeated twice (for a total of four encoding and four retrieval runs). All encoding and retrieval trials were presented for 5 seconds. Participants were provided with practice prior to beginning the task to facilitate learning the two encoding strategies. They were given a total of 8 trials at encoding where participants saw a prompt to use one of the strategies and saw 4 trials then a prompt to use the other strategy and saw another 4 trials. Following encoding practice, they also had 8 trials of retrieval practice with 6 of those trials being targets.

### 2.5 Image Acquisition

Structural and functional images were acquired using a Siemens 3-T scanner equipped with a 12-channel head coil, parallel to the AC–PC plane. Structural images were acquired with a 1650-msec repetition time, a 2.03-msec echo time, a 256-mm field of view, a 2562 matrix, 160 axial slices, and a 1.0-mm slice thickness for each participant. Echoplanar functional images were acquired using a descending acquisition, a 2500-msec repetition time, a 25-msec echo time, a 240-mm field of view, a 802 matrix, a 90° flip angle, and 42 axial slices with a 3.0-mm slice thickness resulting in 3.0-mm isotropic voxels.

### 2.6 Anatomical Data Processing

NIFTI files were preprocessed using the Brain Imaging Data Structure (BIDS; Gorgolewski et al., 2016). Preprocessing was performed using *fMRIPrep* 20.1.1 (Esteban, Markiewicz, Burns, et al., 2022; Esteban, Markiewicz, Goncalves, et al., 2022; RRID:SCR_016216), which is based on *Nipype* 1.5.0 (Gorgolewski et al., 2017; RRID:SCR_002502). A total of 1 T1-weighted (T1w) images were found within the input BIDS dataset. The T1-weighted (T1w) image was corrected for intensity non-uniformity (INU) with N4BiasFieldCorrection (Tustison et al., 2010), distributed with ANTs 2.2.0 (Avants et al., 2008, RRID:SCR_004757), and used as T1w-reference throughout the workflow. The T1w-reference was then skull-stripped with a *Nipype* implementation of the antsBrainExtraction.sh workflow (from ANTs), using OASIS30ANTs as target template. Brain tissue segmentation of cerebrospinal fluid (CSF), white-matter (WM) and gray-matter (GM) was performed on the brain-extracted T1w using fast (FSL 5.0.9, RRID:SCR_002823, Zhang et al., 2001). Brain surfaces were reconstructed using recon-all (FreeSurfer 6.0.1, RRID:SCR_001847, Dale et al., 1999), and the brain mask estimated previously was refined with a custom variation of the method to reconcile ANTs-derived and FreeSurfer-derived segmentations of the cortical gray-matter of Mindboggle (RRID:SCR_002438, Klein et al., 2017). Volume-based spatial normalization to one standard space (MNI152NLin2009cAsym) was performed through nonlinear registration with antsRegistration (ANTs 2.2.0), using brain-extracted versions of both T1w reference and the T1w template. The following template was selected for spatial normalization: *ICBM 152 Nonlinear Asymmetrical template version 2009c* [Fonov et al., (2009), RRID:SCR_008796; TemplateFlow ID: MNI152NLin2009cAsym].

### 2.7 Functional data preprocessing

For each of the 8 BOLD runs found per subject (across all tasks and sessions), the following preprocessing was performed. First, a reference volume and its skull-stripped version were generated using a custom methodology of *fMRIPrep*. Head-motion parameters with respect to the BOLD reference (transformation matrices, and six corresponding rotation and translation parameters) are estimated before any spatiotemporal filtering using mcflirt (FSL 5.0.9, Jenkinson et al., 2002). BOLD runs were slice-time corrected using 3dTshift from AFNI 20160207 (Cox & Hyde, 1997, RRID:SCR_005927). Susceptibility distortion correction (SDC) was omitted. The BOLD reference was then co-registered to the T1w reference using bbregister (FreeSurfer) which implements boundary-based registration (Greve & Fischl, 2009). Co-registration was configured with six degrees of freedom. The BOLD time-series (including slice-timing correction when applied) were resampled onto their original, native space by applying the transforms to correct for head-motion. These resampled BOLD time-series will be referred to as *preprocessed BOLD in original space*, or just *preprocessed BOLD*. The BOLD time-series were resampled into standard space, generating a *preprocessed BOLD run in MNI152NLin2009cAsym space*. First, a reference volume and its skull-stripped version were generated using a custom methodology of *fMRIPrep*. Several confounding time-series were calculated based on the *preprocessed BOLD*: framewise displacement (FD), DVARS and three region-wise global signals. FD was computed using two formulations following Power (absolute sum of relative motions, Power et al., 2014 and Jenkinson (relative root mean square displacement between affines, Jenkinson et al., 2002)). FD and DVARS are calculated for each functional run, both using their implementations in *Nipype* (following the definitions by Power et al., 2014). The three global signals are extracted within the CSF, the WM, and the whole-brain masks. Additionally, a set of physiological regressors were extracted to allow for component-based noise correction (*CompCor*, Behzadi et al., 2007). Principal components are estimated after high-pass filtering the *preprocessed BOLD* time-series (using a discrete cosine filter with 128s cut-off) for the two *CompCor* variants: temporal (tCompCor) and anatomical (aCompCor). tCompCor components are then calculated from the top 5% variable voxels within a mask covering the subcortical regions. This subcortical mask is obtained by heavily eroding the brain mask, which ensures it does not include cortical GM regions. aCompCor components were not utilized in the current set of analyses. For each CompCor decomposition, the *k* components with the largest singular values are retained, such that the retained components’ time series are sufficient to explain 50 percent of variance across the nuisance mask. The remaining components are dropped from consideration. The head-motion estimates calculated in the correction step were also placed within the corresponding confounds file. The confound time series derived from head motion estimates and global signals were expanded with the inclusion of temporal derivatives and quadratic terms for each (Satterthwaite et al., 2013). Frames that exceeded a threshold of 0.5 mm FD or 1.5 standardized DVARS were annotated as motion outliers. All resamplings can be performed with *a single interpolation step* by composing all the pertinent transformations (i.e. head-motion transform matrices, susceptibility distortion correction when available, and co-registrations to anatomical and output spaces). Gridded (volumetric) resamplings were performed using antsApplyTransforms (ANTs), configured with Lanczos interpolation to minimize the smoothing effects of other kernels (Lanczos, 1964). Non-gridded (surface) resamplings were performed using mri_vol2surf (FreeSurfer).

### 2.8 Multivariate Pattern Analyses

To estimate neural activity associated with individual trials, separate GLMs on unsmoothed data were estimated in SPM12 defining one regressor for each trial at encoding and retrieval (172 total for each phase). An additional 6 nuisance regressors were included in each run corresponding to motion. Whole-brain parameter maps were generated for each trial for encoding and retrieval for each participant. In any given parameter map, the value in each voxel represents the regression coefficient for that trial’s regressor in multiple regression containing all other trials in the run and the motion parameters. These beta parameter maps were concatenated across runs and submitted to CoSMoMVPA toolbox (Oosterhof et al., 2016) for pattern classification (Mumford et al., 2012), distinctiveness (Haxby et al., 2001), and reinstatement (Hill et al., 2021) analyses.

#### Multivariate Pattern Analysis (MVPA)

Given our interest in determining which MTL regions discriminated between intra- and inter-item associations, classification analyses were conducted to determine if a classifier was able to discriminate between intra- and inter-item associative targets in our selected ROIs. Separate classification accuracies were computed between the foregoing trial types at both encoding and retrieval using a support vector machine (SVM) classifier with a linear kernel using all voxels within each ROI (Mumford et al., 2012). Training and testing followed an n – 1 cross-validation procedure with three runs used as a training dataset and one run used as testing data. Group-level results were generated from averaging across validation folds from all possible train-data/test-data permutations from the individual participant level. Finally, we tested whether a classifier was significantly able to discriminate neural patterns above chance between the two target types using a one-tailed one-sample t-test for classification accuracy within each ROI. Due to unequal trial numbers in behavioral responses, and the incompatibility with unequal trial numbers in the SVM, the MVPA was conducted only on target trials.

#### Neural Pattern Distinctiveness

Neural distinctiveness has been previously used to examine how distinct neural patterns are from one another in different conditions, and to determine if brain regions are able to discriminate between different conditions or stimuli (J. Haxby et al., 2001; Kriegeskorte et al., 2008; LaRocque et al., 2013). Pattern distinctiveness analyses were conducted to examine the representation of stimuli associated with intra-item and inter-item associative recollected targets. For the purposes of the current analyses, a within condition similarity score was calculated in each participant for each memory phase (e.g. correlation between all beta parameter maps for intra-item associative trials at encoding) and a between condition similarity score in each participant (e.g. One beta parameter map for an intra-item associative trial correlated to all beta parameter maps for inter-associative trials, done for all intra-associative trials and then averaged) (Haxby et al., 2001). Then an overall distinctiveness score was calculated by taking the mean of the intra-within and inter-within similarities and subtracting the between similarity and t-tested against 0 for all ROIs to determine if distinctiveness was above 0. To further examine whether the distinctiveness was driven by within-condition differences, the intra-item’s within similarity and inter-item’s within similarity was directly compared using a paired t-test within each ROI. This process was repeated for both encoding and retrieval trials.

#### Category-level Neural Pattern Reinstatement

In parallel with Hill and colleagues (2021), we used category-level reinstatement to determine whether reinstatement differed between associative conditions at the category-level. These analyses were run to examine how different associative recollected target information is reinstated from encoding to retrieval, which may help determine the mechanism by which intra-item associations function neurally compared to inter-item associations. First, we calculated a within condition similarity value in each participant in each ROI (e.g. one beta parameter map for an intra-item associative trial at encoding correlated to all beta parameter maps for intra-item associative trials at retrieval and paralleled for the inter-associative condition). Then we calculated a between condition similarity value in each participant in each ROI (e.g. one beta parameter map for an intra-item associative trial at encoding correlated to all beta parameter maps for inter trials at retrieval and vice versa for inter-associative condition). Next, we calculated an overall reinstatement score by taking the mean of intra- and inter-item associative within encoding retrieval similarity and subtracting the mean of the between intra- and inter-item associative encoding retrieval similarity and used a one-tailed, one-sample t-test against 0 for all ROIs to determine if the reinstatement was above chance (*µ*_*within-category*_*-µ*_*between-category*_). Next, we calculated an intra-item associative category-reinstatement value by subtracting the mean within intra-item associative value from the mean between category value and the same was done for the inter condition. Finally, to directly compare the encoding conditions, the separate intra-and inter-item associative category-level reinstatement were compared via a paired t-test (See figure 2).

**Figure 2.**
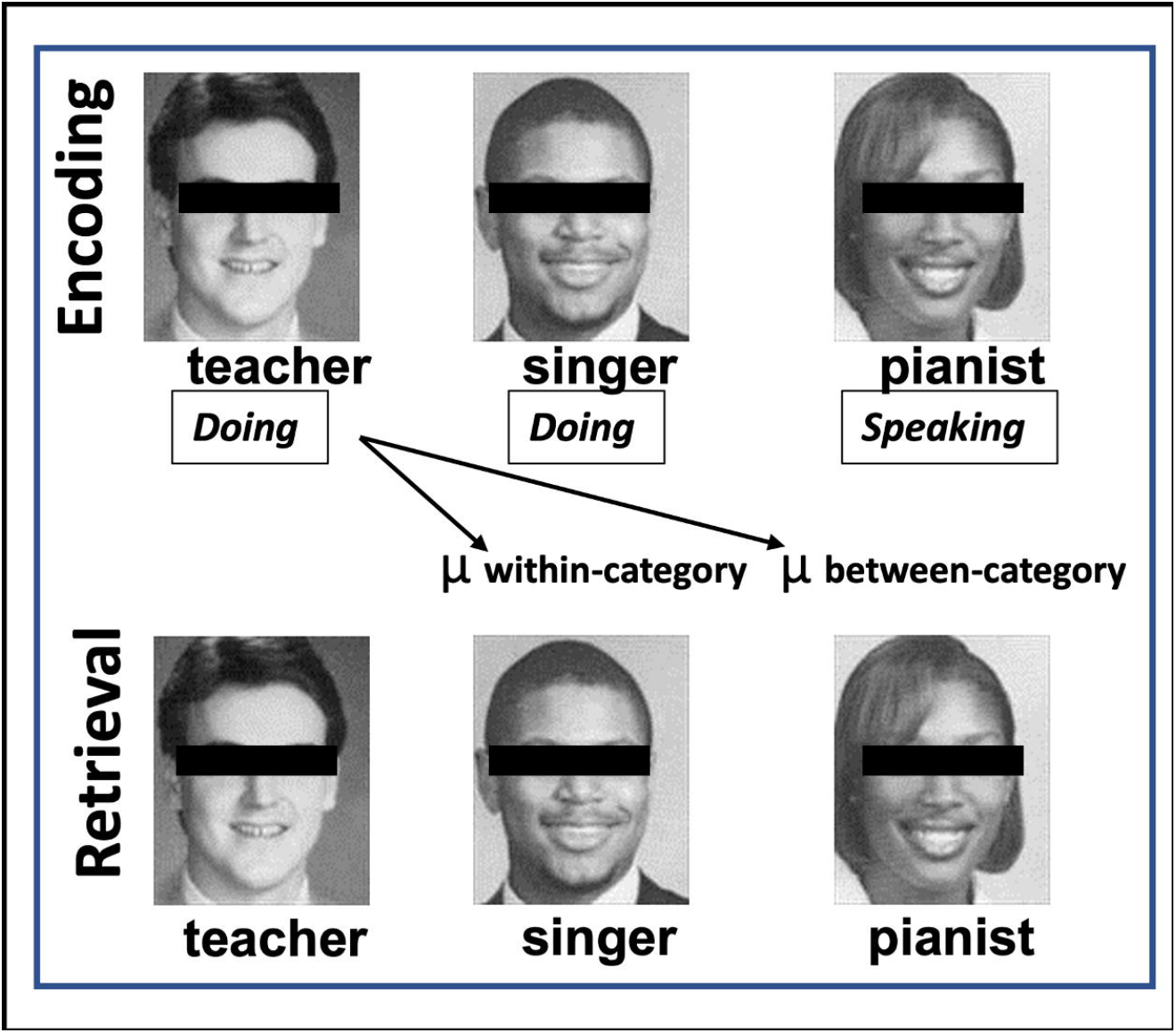
Example showing Category-Level Reinstatement. Category-Level reinstatement was calculated by the mean correlation of within category (i.e., doing condition) minus the mean correlation of the between category items (i.e., speaking condition). Category-Level Reinstatement = µ_within-category_ - µ_between-category_

## 3. Results

### Behavioral Results

A series of paired t-tests were run between intra- and inter-item associative trials for each of the following: recollected hits, correct rejections (CR), and reaction times (RT). Comparing the recollected hit rate for intra- and inter-item associative trials showed significant differences (intra-item associations: M=0.58 SD=0.13, inter-item associations: M=0.47 SD=0.15; *t*(27)=5.83, *p*<.001, 95% CI [0.06, 0.013]). Comparing correct rejections between intra- and inter-item associations showed no significance (intra-item associations: M=0.59 SD=0.15, inter-item associations: M=0.56 SD=0.16; *t*(27)=1.36, *p*=.19, 95% CI[-0.02, 0.08]).

### Classification Results

To examine whether classifiers were able to significantly discriminate between our two target conditions, two multivoxel pattern analyses were run. The first was to classify all intra- and inter-item associative targets at encoding, and the second to classify all intra- and inter-item associative targets at retrieval. Comparing classification of intra- and inter-item associative targets at encoding against chance (50%) showed no above chance significance within any ROI (HC: *t*(27)=0.19, *p*=.85, 95% CI[0.48, 0.53]; PHC: *t*(27)=-0.41, *p*=.69, 95% CI[0.47, 0.52]; PrC: *t*(27)=1.08, *p*=.29, 95% CI[0.49, 0.55]). Comparing classification of intra- and inter-item associative targets at retrieval against chance (50%) revealed significance in the HC, such that classifier accuracy was significantly above chance, (*t*(27)= 2.56, *p*=.016, 95% CI[0.51, 0.55]). The PHC and PrC did not show above chance classifier accuracy (PHC: *t*(27)=0.26, *p*=.80, 95% CI[0.48,0.52]; PrC: *t*(27)=0.92, *p*=.37, 95% CI[0.49,0.54]).

### Encoding Distinctiveness

In order to examine neural discriminability between two successful encoding conditions, a neural distinctiveness calculation was conducted for intra- and inter-item association recollected targets (within category similarity minus between category similarity). The distinctiveness in both conditions was then compared to determine if one condition showed higher sensitivity in a given region compared to the other condition. At encoding, overall distinctiveness of recollected targets, collapsed across condition, was significantly greater than 0 within all ROIs [HC: *t*(27)=9.72, *p*<.001; PHC: *t*(27)=6.88, *p*<.001; PrC: *t*(27)=13.11, *p*<.001]. When breaking down the distinctiveness of recollection by condition we found that both conditions were significantly greater than 0 in all ROIs [intra-item associative targets: HC: *t*(27)=6.40, *p*<.001; PHC: *t*(27)=5.40, *p*<.001; PrC: *t*(27)=5.47, *p*<.001; inter-item associative targets: HC: *t*(27)=7.63, *p*<.001; PHC: *t*(27)=4.92, *p*<.001; PrC: *t*(27)=5.35, *p*<.001;]. A direct comparison of within-condition similarity between conditions found no significant differences in any ROI [HC: *t*(27)=0.29, *p=*.78, 95% CI[-0.01, 0.01]; PHC: *t*(27)=0.15, *p=*.88, 95% CI[-0.02, 0.02]; PrC: *t*(27)=-0.49, *p*=.63, 95% CI[-0.02, 0.01]]. (See figure 3).

**Figure 3.**
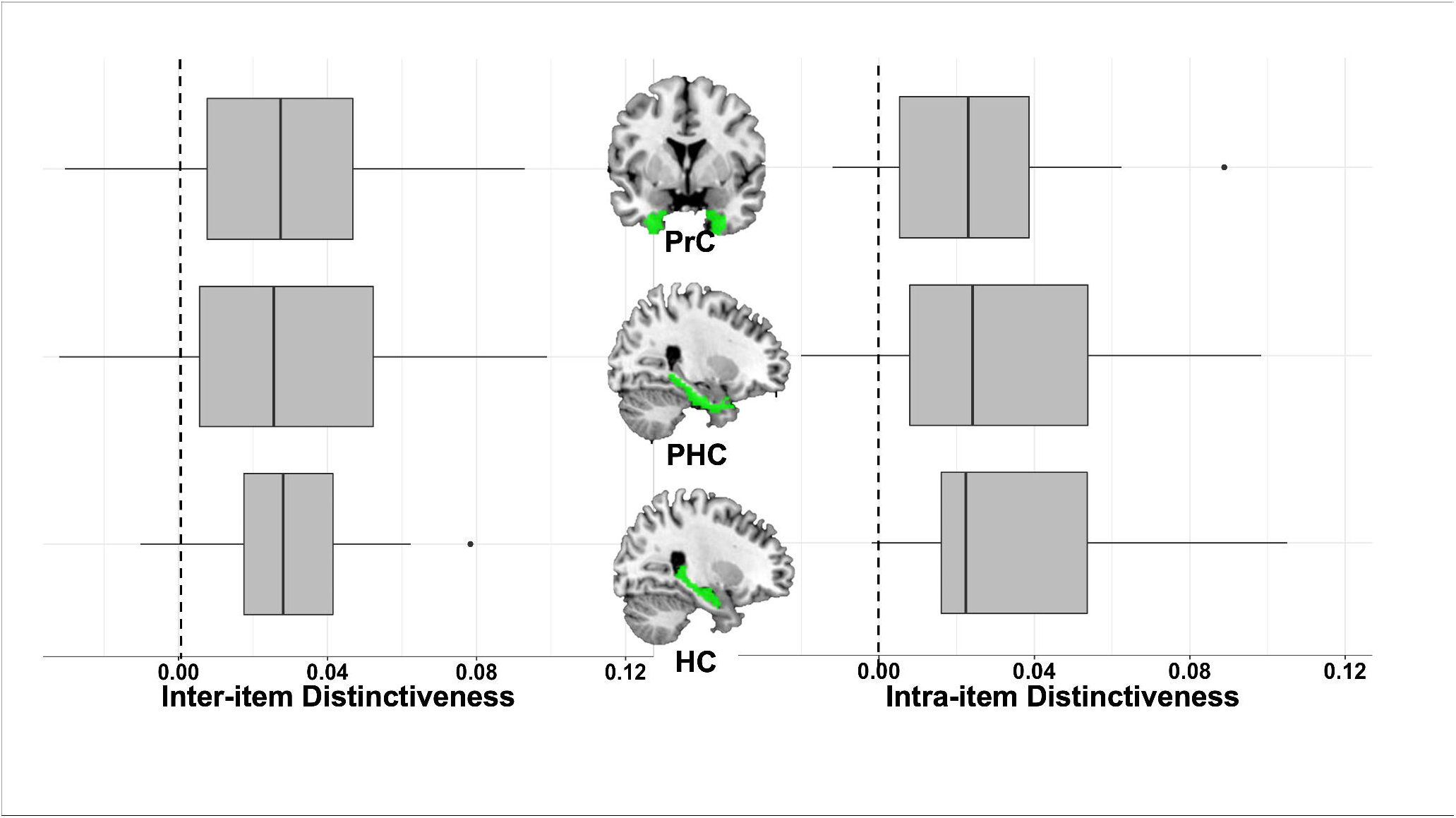
Neural distinctiveness during encoding in the perirhinal cortex (PrC), parahippocampal cortex (PHC), and the hippocampus (HC) across the two encoding conditions. All above neural distinctiveness scores were significantly different than 0; all *ps* < 0.001.

### Retrieval Distinctiveness

In order to examine neural discriminability between two successful retrieval conditions, a neural distinctiveness calculation was conducted for intra- and inter-item association recollected targets (within category similarity minus between category similarity). The distinctiveness in both conditions was then compared to determine if one condition showed higher sensitivity in a given region compared to the other condition. At retrieval, overall distinctiveness of recollected targets, collapsed across condition, was not significantly greater than 0 within any ROIs [HC: *t*(27)=-0.53, *p*=.70; PHC: *t*(27)=-0.48, *p*=.68; PrC: *t*(27)=0.66, *p*=.26]. Similarly, when breaking down the distinctiveness of recollection by condition we found the distinctiveness of recollected intra-item associations [HC: *t*(27)=-0.20, *p*=.58; PHC: *t*(27)=-1.33, *p*=.90; PrC: *t*(27)=-0.69, *p*=.75] and inter-item associations [HC: *t*(27)=-0.42, *p*=.66; PHC: *t*(27)=0.71, *p*=.24; PrC: *t*(27)=1.54, *p*=.07] were not significantly greater than 0 within any ROI.

### Neural reinstatement

To examine how different associative recollected target information is reinstated from encoding to retrieval, neural reinstatement was calculated for intra- and inter-item associations and then compared to one another to determine if reinstatement was greater in one condition compared to the other. Category level neural reinstatement was conducted on recollected targets with an overall category level, collapsed across conditions, and two others separated by condition. Overall category level reinstatement compared against 0, reveals that there is significant reinstatement of recollected targets regardless of trial type, within the HC [*t*(27)=3.87, *p*<.001] and PHC [*t*(27)=4.54, *p*<.001]. However, the PrC did not show significant reinstatement across trial types [*t*(27)=-5.38, *p*=.99]. Intra-item associative category level reinstatement t-tests compared against 0 revealed that the HC [*t*(27)=3.25, *p*<.01] and PHC [*t*(27)=3.89, *p*<.001] again show significant reinstatement from encoding to retrieval. The PrC did not show significant reinstatement of recollected intra-item associative targets [*t*(27)=-3.69, *p*=.99]. Inter-item category level reinstatement t-tests compared against 0 revealed that the HC [*t*(27)=4.10, *p*<.001] and PHC [*t*(27)=4.88, *p*<.001] show significant reinstatement from encoding to retrieval. The PrC did not show significant reinstatement of recollected inter-item associative targets [*t*(27)=-6.44 *p*=.99]. To directly compare reinstatement between intra- and inter-item associative recollected targets, we compared the two trial types in the regions of significance. The results revealed a significant difference between reinstatement of intra- and inter-item associative recollected targets in the HC [*t*(27)=2.81, *p*<.01], such that recollected inter-item associative targets were reinstated from encoding to retrieval to a greater extent than recollected intra-item associative targets. The PHC however did not show any significant differences between intra- and inter-item associative trials [*t*(27)=1.90, *p*=.07]. (See figure 4).

**Figure 4.**
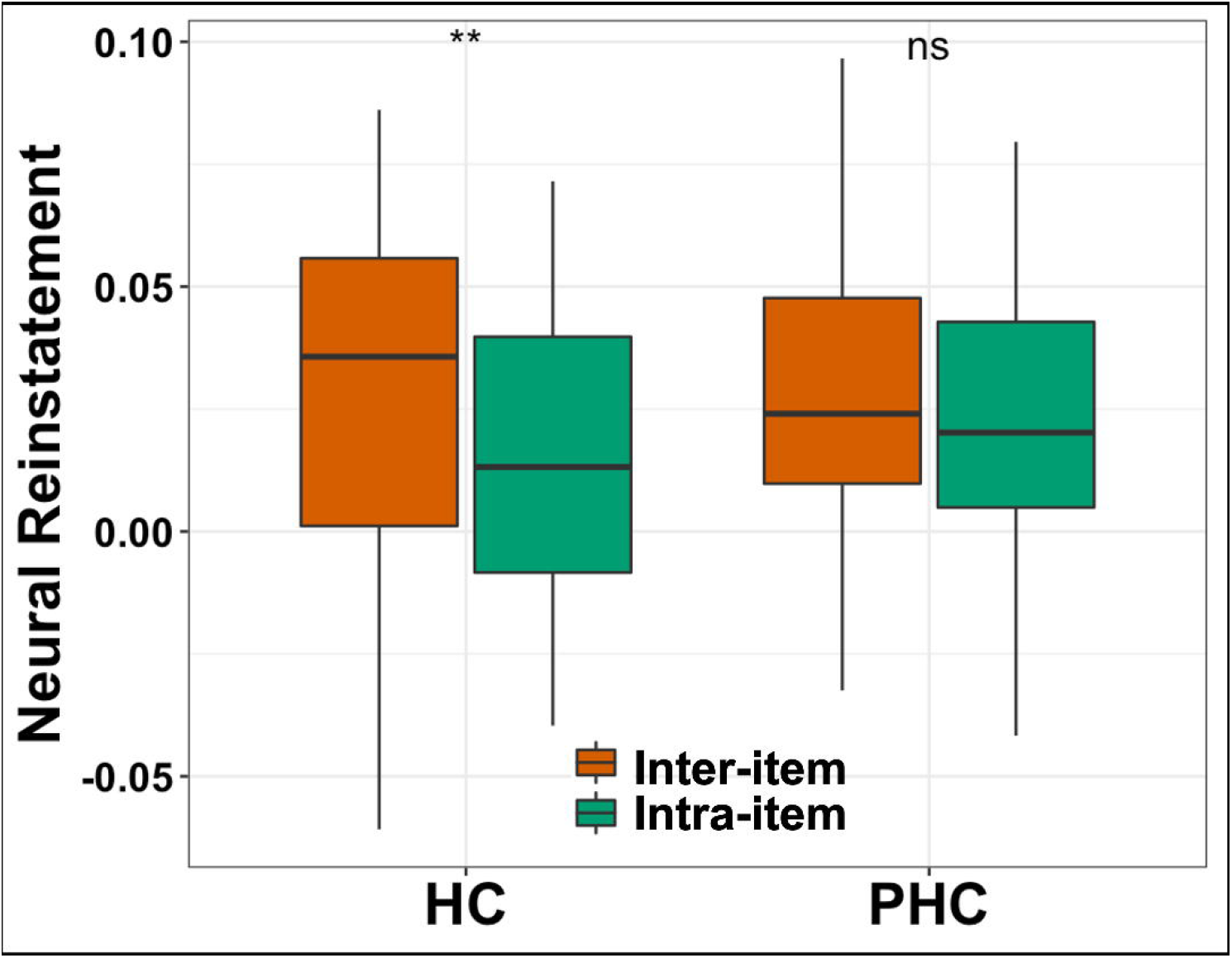
Neural reinstatement (within-between) of doing and speaking trials, as t-tested against one another. **: *p*<.01.

## 4. Discussion

The goal of the current study was to examine the underlying neural mechanisms of intra-item compared to inter-item associations, induced through the manipulation of encoding instructions. Behavioral results demonstrate that participants were more successful at recollecting face-occupation associations that were encoded using intra-item binding compared to those encoded using inter-item binding. These findings replicate those observed by Overman & Stephens (2013) who also showed that associative memory was enhanced with an encoding task that emphasized intra-item binding. It is hypothesized that such integrative encoding operates by promoting unitization of items in memory (Bastin et al., 2013; Delhaye et al., 2014; Giovanello et al., 2006; Parks & Yonelinas, 2015) and creating a holistic representation through internalizing the association of two items. This is different than inter-item binding which forms an external link between two items (Graf & Schacter, 1989; Park & Rugg, 2011). Given the foregoing differences in associative recollection across encoding conditions, we sought to identify the neural basis underscoring this behavioral advantage of intra-item compared to inter-item binding. We hypothesized that this behavioral advantage would stem from higher neural distinctiveness of intra-item pairs, and greater reinstatement between encoding and retrieval of subsequent intra-item pairs, compared to inter-item pairs.

### Multivariate Classification

Previous work focusing on intra-item associations, in the form of unitization, finds that inter-item processing occurs in the hippocampus, while intra-item binding is processed by the perirhinal cortex, thus the two types of memory are thought to utilize distinct neural processes (Eichenbaum et al., 1994; Haskins et al., 2008; Jäger et al., 2006; Moses & Ryan, 2006; Staresina & Davachi, 2010). While this has been investigated from a univariate perspective, it was unclear if distributed patterns of neural activity related to intra- or inter-item processing were discriminable. Multivoxel pattern classification allowed us to investigate whether intra- and inter-item associations are discriminable from one another at encoding and retrieval. While no differences were found within MTL subregions during encoding, we found the two types of associations were discriminable during retrieval within the hippocampus. This suggests that the hippocampus specifically, compared to other MTL regions, is sensitive to different types of associative pairings at retrieval, whether that be a more wholistically encoded association, or a more loosely bound association (Diana et al., 2008; Ranganath, 2010; Staresina & Davachi, 2010). While previous work typically implicates the PrC in supporting intra-item pairings (Haskins et al., 2008; Jäger et al., 2006; Staresina & Davachi, 2010), an absence of discriminability within this region may suggest that for more similar stimuli (e.g. face-job pairings differing only by strategy instructions) the hippocampus is necessary to parcellate between more fine-grained differences in the retrieval the bound information.

### Neural Distinctiveness

We next calculated the neural distinctiveness (Haxby et al., 2001) of our associative conditions to investigate whether neural patterns of successfully recollected intra- and inter-item associations were unique from one another at each memory phase (i.e. encoding and retrieval). Results showed that at encoding, yet not retrieval, neural patterns related to intra- and inter-item associations were discriminable from one another within all MTL subregions, including the PrC, PHC, and HC. However, there were no differences in the within-condition pattern similarity, or the neural consistency, of this metric across associative categories, suggesting that both association types may show similar degrees of reliability within their neural representations (Simmonite & Polk, 2022). This suggests that while utilizing similar MTL regions, information in each condition is encoded distinctly within these regions in a manner that allows for subsequent successful recollection. However, at retrieval, where there is no significant distinctiveness in any ROI, these different memory traces may be processed and retrieved successfully in a generalized manner that lacks the distinct pattern differences found during the initial processing of the associative pair.

Classification of targets at retrieval, and distinctiveness of subsequentially recollected trials at encoding suggest that processes when encoding associative pairs are critical to their subsequent successful recollection compared to the retrieval of targets irrespective of behavior. While these discrimination processes are evident for both intra- and inter-item associations, these results are consistent with prior work featuring univariate contrasts and event-related potentials in associative memory that suggest intra-item associations are bound into a unitized representation during encoding, and that differences in the strength of the bond results in intra-item associations success over and above that of inter-item associations at retrieval (Haskins et al., 2008; Jäger et al., 2006; Staresina & Davachi, 2010). Specifically for intra-item associations, the PrC has been implicated in the encoding processes of intra-item or unitized associations, while inter-item associations tend to be more reliant on HC and PHC regions (Allen et al., 2014; Dennis, Johnson, et al., 2014; Haskins et al., 2008; Piekema et al., 2010; Staresina & Davachi, 2010; Yonelinas et al., 2001). However, the current results suggest that both intra- and inter-item associations utilize all MTL subregions during the encoding process. The current results also suggest that binding of both intra- and inter-item associations is supported at encoding by the PrC as well as other MTL regions. These findings support the notion that intra- and inter-item associations are distinct processes, while utilizing similar regions to successfully encode associative information. This may again be in the nature of the stimuli and the encoding instructions, such that all MTL regions may be critical in distinguishing between the two types of stimuli in order to recollect more fine-grained differences between similar stimuli (Bussey et al., 2002; Moss et al., 2005; Murray & Richmond, 2001).

### Neural Reinstatement

While the foregoing results suggest that intra and inter-item associations are discriminable from one another when examining neural patterns associated with successful recollection of the associated pairs, it was also of interest to investigate how information related to intra- and inter-item association is recapitulated across study phases. The results from our neural reinstatement analysis suggest that neural patterns underlying the processing of successfully recollected intra- and inter-item associations are reinstated within the HC and PHC, and are reinstated distinctly from one another in these regions. inter-item associations Specifically, we found that inter-item associations exhibited greater reinstatement in the HC compared to intra-item associations. While this finding was contradictory to our hypothesis, the current results are suggestive of the idea that intra-item associations promote more efficient associations (Parks & Yonelinas, 2015; Quamme et al., 2007) and thus, should require less neural resources to be reinstated cross memory phases. The current reinstatement results also suggest that greater reinstatement in inter-item associations may indicate a compensatory process (Schmidt et al., 2019), such that neural representations must be reinstated to a greater degree in order to recollect inter-item associations to the same extent that intra-item trials are successfully recollected. While the behavioral metrics did not correlate with neural reinstatement in any ROI, behavioral results did show that inter-item associations are recollected less often than intra-item associations which, in itself, suggests that there is some inherent difference in the way in which the two associative types are processed neurally. This also suggests that even younger adults may use some compensatory mechanisms while recapitulating associative information that is relatively difficult to bind. This is especially critical, such that intra-item pairings are typically found to be of most benefit to older adult populations (Ahmad & Hockley, 2014; Bastin et al., 2013; Delhaye & Bastin, 2018).

Taken together with the above neural discriminability findings, our results suggest that the neural reinstatement of successfully recollected intra- and inter-item associations is driven by differences at encoding rather than retrieval (Staresina & Davachi, 2010; Tu & Diana, 2021). This is consistent with behavioral and early neuroimaging work with intra-item associations that suggest that these associations allow for a more efficient binding of associative information at encoding compared to inter-item associations, thus accounting for the behavioral advantage (Ahmad & Hockley, 2014; Bastin et al., 2013; Delhaye & Bastin, 2018; Jäger et al., 2006; Parks & Yonelinas, 2015; Staresina & Davachi, 2010). The current set of results adds to this prior work, providing evidence that not only do these regions exhibit altered BOLD recruitment when processing these associations, but that neural patterns are discriminated and reinstated differentially as well.

## 5. Limitations/Future Directions

While the current findings highlight the role of MTL subregions in recollection across intra- and inter-item associative memory, we were unable to look at other retrieval-based processes such as familiarity due to low trial counts. This could be a fruitful avenue for future research, especially given the link between unitization and familiarity in memory and the role the PrC may play in both processes. Additionally, we cannot be certain that the intra-item condition is truly being bound as a single unit, only that behavioral differences across encoding conditions suggest stronger memory for that trial type. Without a single item condition as a comparison, the intra-item association can only be concluded as being different from the inter-item associations. Future work should aim to identify whether these differences in intra- and inter-item associations are due to unitization processing and whether they differ as we age. Specifically, it should be determined if the compensatory process described earlier is also observable in older adult populations.

## 6. Conclusions

Taken together, the results suggest neural patterns relevant to intra- and inter-item associations are discriminable and are reinstated differentially between memory phases. While intra-item associations lead to more successful memory compared to inter-item associations behaviorally, the neural mechanisms underlying these two types of memories may be of a compensatory nature. With respect to neural discriminability of successfully recollected intra- and inter-item associations, these associations appear to utilize similar MTL regions to successfully encode and discriminate the associative information. Additionally, inter-item associations, in order to perform as successfully as intra-item associations, may utilize compensatory reinstatement processes to a greater extent compared to intra-item associations and that this compensatory process is discernable in young adult populations.

## Acknowledgements

We would like to thank Harini Babu, Kayla McGraw, Valerie Goodwin, Chloe Hultman and Min Sung Seo for helping with data collection and analysis. This paper was supported by the National Institutes of Health under grant R15 AG052903 awarded to A.A.O and N.A.D.

## Availability of Data and Materials

The datasets generated and analyzed during the current study are available from the corresponding author on reasonable request. None of the experiments were preregistered.

## Footnotes

1 Nine participants completed this version of task before it was changed to alternate between instructions after 18 trials (N=19), and prompt lasting 5 seconds. There were no repetitions of strategy in each run (i.e. subjects saw one doing and one speaking strategy block per run). This change was made in response to feedback from older adults. There were no significant differences in any result between versions.

2 Due to an error in programming, a subset of lures were recombined between (rather than within) condition. All between condition lures were removed from behavioral and imaging analyses.

